# Profiling of HeLa nucleoplasmic and nucleolar RNAs by Halo-seq proximity labeling

**DOI:** 10.64898/2026.01.29.702496

**Authors:** Timo Ahola, Piia Takabe, Riku Salomäki, Päivi Lillsunde, Mohamed Bahrudeen, Donald Wlodkowic, Jan Kaslin, Alexander Efimov, Minna-Liisa Änkö

## Abstract

Subcellular RNA localization is a key regulator of gene expression, but transcriptome wide characterization of RNAs enriched in specific cellular compartments has been hindered by methodological limitations in throughput and spatial resolution. Halo-seq is an RNA proximity-labeling approach that enables the extraction of RNAs located near virtually any Halo-tagged bait protein. However, its broader application has been restricted by the limited availability of the required Halo ligand. Here we present a Halo-seq dataset profiling the nucleoplasmic and nucleolar transcriptomes of HeLa cells. Through experimental and computational validation, we demonstrate that our optimized Halo-seq protocol achieves high spatial specificity, enabling robust mapping of compartment-enriched RNA populations.

## Background and summary

Across species and cell types, RNAs are not evenly distributed within cells, and this spatial asymmetry in transcriptomes plays a crucial role in regulating cellular functions ^1^. Membrane-bound compartments such as the nucleus restrict RNA movement, giving rise to distinct subcellular transcriptomes. Even within these compartments, RNAs exhibit precise localization patterns. For example, nuclear RNAs can accumulate in specific nuclear organelles, whereas cytosolic RNAs may be targeted to the cell membrane ^2^. Because RNA localization influences translation, processing, and stability, alterations in RNA positioning can profoundly impact gene expression and, consequently, cellular function ^3^.

RNA localization can be studied at high resolution using fluorescent *in situ* hybridization (FISH) -based approaches. However, FISH remains limited in throughput because each investigated RNA requires a predesigned probe set, and the number of available detection channels further constrains scalability ^4–6^. Newer multiplexing FISH methods, like MERFISH and seqFISH, partially overcome these limitations, but still fall short of achieving transcriptome-wide coverage and require extensive optimization ^7,8^.

Biochemical and mechanical fractionation strategies combined with RNA-sequencing (RNA-seq) offer another means to profile subcellular RNA content ^9,10^. These approaches are limited in spatial resolution because they are restricted to compartments that can be biochemically separated, such as the nucleus or plasma membrane. Recently, fluorescence-based sorting approaches have been applied to isolate specific cellular condensates ^11–13^.

Proximity-labeling (PL) methods use bait proteins to isolate RNAs located in their immediate vicinity. The most widely used PL RNA-seq strategy, APEX-seq, employs engineered soybean ascorbate peroxidase (APEX2) -tagged bait proteins to generate biotin-phenol radicals that covalently attach to nearby RNAs ^14,15^. Halo-seq is an alternative PL approach designed to increase the efficiency and specificity of RNA labeling by using a non-enzymatic light-dependent labeling reaction ^16^. In Halo-seq, RNAs are labeled through Halo-tagged bait proteins bound to a dibromofluorescein (DBF) HaloTag ligand. Upon green light exposure, the HaloTag-conjugated DBF produces reactive oxygen species (ROS) that oxidise guanines in proximal RNAs to 8-oxoguanines. Subsequent treatment of cells with the nucleophile propargylamine (PA) enables covalent attachment to the oxidated guanines. A copper-catalysed Azide-Alkyne Cycloaddition (CuAAC) ‘click’ reaction is then used to conjugate a biotin-azide to PA, allowing enrichment of the biotinylated RNAs using streptavidin beads to generate a Halo-seq pulldown library ^17^. The Halo-seq labeling workflow is presented in **Figure 1**.

**Figure 1.**
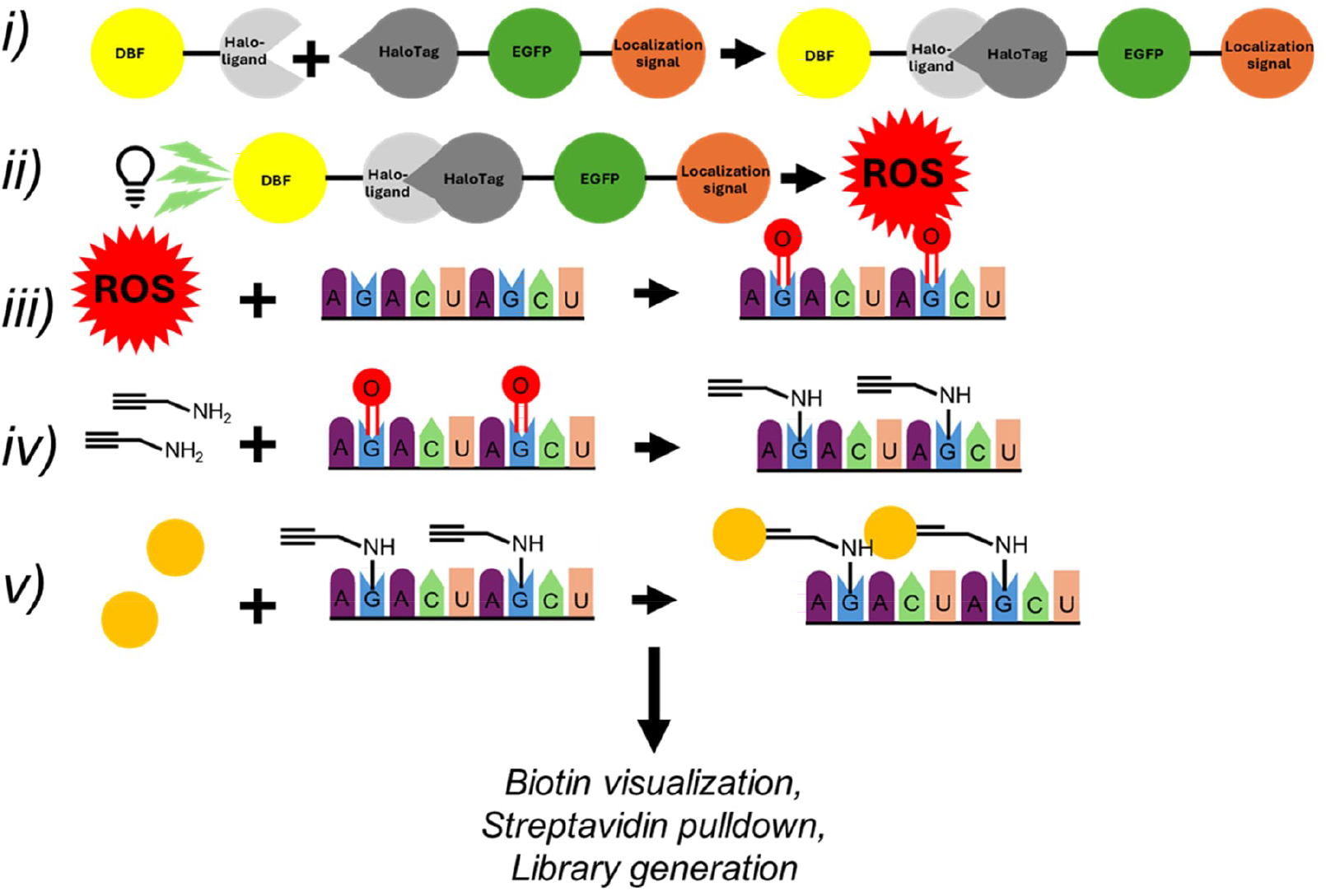
A schematic of Halo-seq RNA labelling. i) The conjugation of DBF-ligand to HaloTagged protein. ii) Light activated ROS generation by DBF. iii) Oxidation of nearby guanines. iv) Covalent attachment of PA through nucleophilic attack. v) CuAAC -mediated attachment of biotin-azide. After biotinylation the labelled RNAs can be visualized, enriched using streptavidin beads, and sequenced.

Applications of Halo-seq have remained limited, likely because the required DBF-ligand is not commercially available and must be synthesized in-house. We optimized the previously described synthesis protocol to obtain highly pure DBF ligand with improved yield (**Supplemental Text 1**) ^18^. Using our DBP ligand, we generated a Halo-seq dataset emplying two Halo-tagged bait proteins, HaloTag-EGFP-Nuclear Localization Signal (Halo-NLS) and HaloTag-EGFP-Fibrillarin (Halo-FBL). Halo-NLS localizes to the nucleoplasm and labels nucleoplasmic RNAs, whereas Halo-FBL taregts the nucleolus and labels nucleolar RNAs. We established HeLa cell lines stably expressing each bait protein at moderate levels and performed Halo-seq in three independent biological replicates. This dataset provides detailed insights into differences between the nucleoplasmic and nucleolar transcriptomes and serves as an independent demonstration of both DBF-ligand synthesis and optimized Halo-seq application.

## Methods

### Generation of cell lines expressing Halo-fusion proteins

The transfer vectors carrying the Halo-fusion constructs (HaloTag-EGFP-NLS and Halo-EGFP-Fibrillarin) were co-transfected with lentiviral packaging vectors, psPAX2 (Addgene #12260) and pMD2G (Addgene #12259), into HEK293-cells. The lentiviral particles were collected 24 and 36h post-transfection and concentrated 70-75-fold using 10 kDa Amicon Ultra Centrifugal Filters (Sigma-Aldrich, USA, UFC901024). A 1:500 dilution of each concentrated vector was used for infection, resulting in MOI<10. The success of transduction was verified by visualizing the EGFP-expression. The generated stable cell lines were sorted and single-cell cloned at the Tampere University Flow Cytometry Facility. For both cell lines, single-cell clones that expressed the fusion protein at a moderate level were selected.

### Fluorescent visualization of biotinylated RNA

The synthesis of the DBF-ligand is presented at **Supplemental text 1**.

The success of Halo-proximal RNA tagging was validated by tagging the alkynylated Halo-proximal RNA with a fluorescent Cy3-azide (Click Chemistry Tools, 1178). Cells grown on coverslips were washed with DPBS and incubated with 0.5 µM DBF-ligand in HBSS (Thermo Fisher Scientific, 14025092) for 15 minutes in culture conditions (37 °C, 5 % CO2). The cells were washed twice with culture media for 10 min followed by incubation with 1 mM propargylamine (Thermo Fisher Scientific, 131460050) in HBSS for 5 min in culture conditions. Cells were exposed to green light at 1700 lux for 5 min at culture conditions. Illumination was performed using a custom-made light source with two led bulb covered plates, between which the culture dish was sandwiched in the incubator. After illumination, the cells were washed twice with DPBS at room temperature (RT) and fixed with 4 % paraformaldehyde (PFA) in DPBS for 10 minutes at RT followed by two washes with DPBS.

To perform the click-reaction, the cells were blocked in 1 mg/ml BSA, 75nM NaCl, 0.025% (m/V) sodium azide, 0.1 % (V/V) Triton-X for 30 minutes at RT followed by washing with DPBS. The click-reaction was performed with 10 mM Tris Ph 7.0, 10 μM CY3-picolyl azide (Click Chemistry Tools, 1178), 10 mM freshly made sodium l-ascorbate, 1 mM THPTA (Sigma-Aldrich, 762342), and 100 μM CuSO_4_. The reaction was incubated for 1 hour at 37 °C in the dark. After incubation, the samples were counterstained with 300 nM DAPI, mounted and imaged with a Nikon Eclipse Ti2 inverted microscope, using a 60X oil-immersion objective. Images were processed with FIJI by linear contrast and brightness adjustments for all channels ^19^.

### Extraction and biotinylation of alkynylated RNA

One confluent Ø15 cm dish of Halo-NLS expressing cells or two dishes of Halo-FBL cells were washed with DPBS followed by 15 minutes incubation with 0.5 µM DBF-ligand in HBSS at culture conditions. Cells were washed twice with culture media for 5 minutes at culture conditions followed by a 5-minute incubation with 1 mM PA in HBSS and illumination with 1700 lux green light at for 5 minutes at culture conditions. After illumination, the cells were lysed with TRIzol® (Thermo Fisher Scientific, 15596018) and the samples were stored at -80°C. Isolated RNA was DNase-treated with TURBO DNase (2 U/µl) (Thermo Fisher Scientific, AM2238) according to manufacturer’s instructions. After DNase treatment, RNA was recovered using Acid-Phenol:Chloroform (Thermo Fisher Scientific, AM9720) and reconstituted in nuclease-free water.

RNA was diluted to a concentration of 300 ng/µl and 0.5X sample volumes of click reaction solution (10 mM Tris Ph 7.0, 2 mM Biotin-picolyl azide (Sigma-Aldrich, 900912), 10 mM freshly made sodium l-ascorbate, 1 mM THPTA, and 100 μM CuSO_4_) was added to the diluted RNA. The samples were divided into 50 µl aliquots and incubated for 30 minutes at 25°C in the dark. The samples were transferred to ice to stop the reaction. RNA was recovered using Monarch RNA cleanup Kit (New England Biolabs, T2050L). Five µg of RNA was stored as input sample for RNA-sequencing.

### Streptavidin pulldown

Biotinylated RNA was enriched using Streptavidin Magnetic Beads (New England Biolabs, S1420S). The beads were washed three times with Binding & Washing (BW)-buffer (10 mM Tris Ph 7.5, 1 M NaCl, 1 mM EDTA, 0.2% RNaseOUT). The beads were collected with DynaMag-2 magnet (Thermo Fisher Scientific, 12321D) for one minute. One hundred twenty µg of biotinylated RNA in BW was incubated with the washed beads for 2 h in 4 °C. After incubation the beads were again washed thrice with the BW-buffer.

After the pulldown, the beads were resuspended in 50 µl of DPBS. The biotin-streptavidin bond was broken by adding 300 µl of TRIzol® to the samples followed by 10 minutes incubation at 37°C. The beads were then separated using the DynaMag-2 and RNA was then extracted.

### RNA-sequencing

The integrity of the samples was evaluated using Agilent Tape station, after which ribo-depleted libraries were generated and sequenced at Novogene (150 bp paired-end reads).

### Sequencing quality control and read mapping

Quality control and preprocessing of RNA-seq reads was performed using fastp ^20^. The reads passing the quality control were mapped using Salmon (version 1.10.2) to an index created from a human transcriptome reference obtained from GENCODE version 46 ^21,22^. Transcript abundances were converted to gene abundances using tximport^23^. Differential gene abundance between pulldown and input samples was analysed using DESeq2 ^24^.

### Manual clustering

Gene clusters were manually curated based on DEseq2 results. Clusters were constructed based on enrichment and lack of detectable enrichment in Halo-seq pulldowns. A gene was considered enriched if it received a padj < 0.05 and log_2_ fold change (LFC) > 0.5 in a pulldown. A lack of detectable enrichment of a gene in a pulldown was assigned if a gene did not receive padj < 0.05 and LFC > 0. Cluster 1 contained genes enriched in both pulldowns. Cluster 2 contained genes enriched in FBL with no detectable enrichment in NLS. Cluster 3 contained genes enriched in NLS with no detectable enrichment in FBL. Cluster 4 contained genes depleted in both pulldowns, meaning they received a padj < 0.05 and LFC < -0.5.

### Gene ontology analysis

Gene ontology analysis was performed for the generated cluster with the clusterprofiler R package ^25^. Only protein coding genes in each cluster were included in the analysis and the universe was set to all protein coding genes expressed in the Halo-seq samples.

### Technical validation

Spatial accuracy of RNA alkynylation was assessed by in-cell Cy3-azide labeling of the alkynylated RNA. As expected, the Halo-fusion proteins and the corresponding labeled RNA colocalized (**Figure 2a**). Halo-NLS protein and its proximally labeled RNA were detected throughout the nucleoplasm, whereas Halo-FBL protein and the associated labeled RNA were confined to the nucleolus. Streptavidin pulldown yields differed between the two cell lines, reflecting the broader distrubtion of Halo-NLS compared with the more restricted localization of Halo-FBL (**Figure 2b**). NLS-Halo samples yielded 1-1.75% of input RNA, whereas Halo-FBL only 0.2-0.48%. because of the lower recovery in Halo-FBL RNA samples, two pulldown reactions were pooled for each biological replicate to obtain sufficient RNA for sequencing library preparation. RNA integrity was assessed using the Agilent Bioanalyzer 2100 prior to sequencing, and all samples displayed ribosomal integrity numbers (RIN) between 7.1 - 8.4 indicating RNA-quality suitable for sequencing.

**Figure 2.**
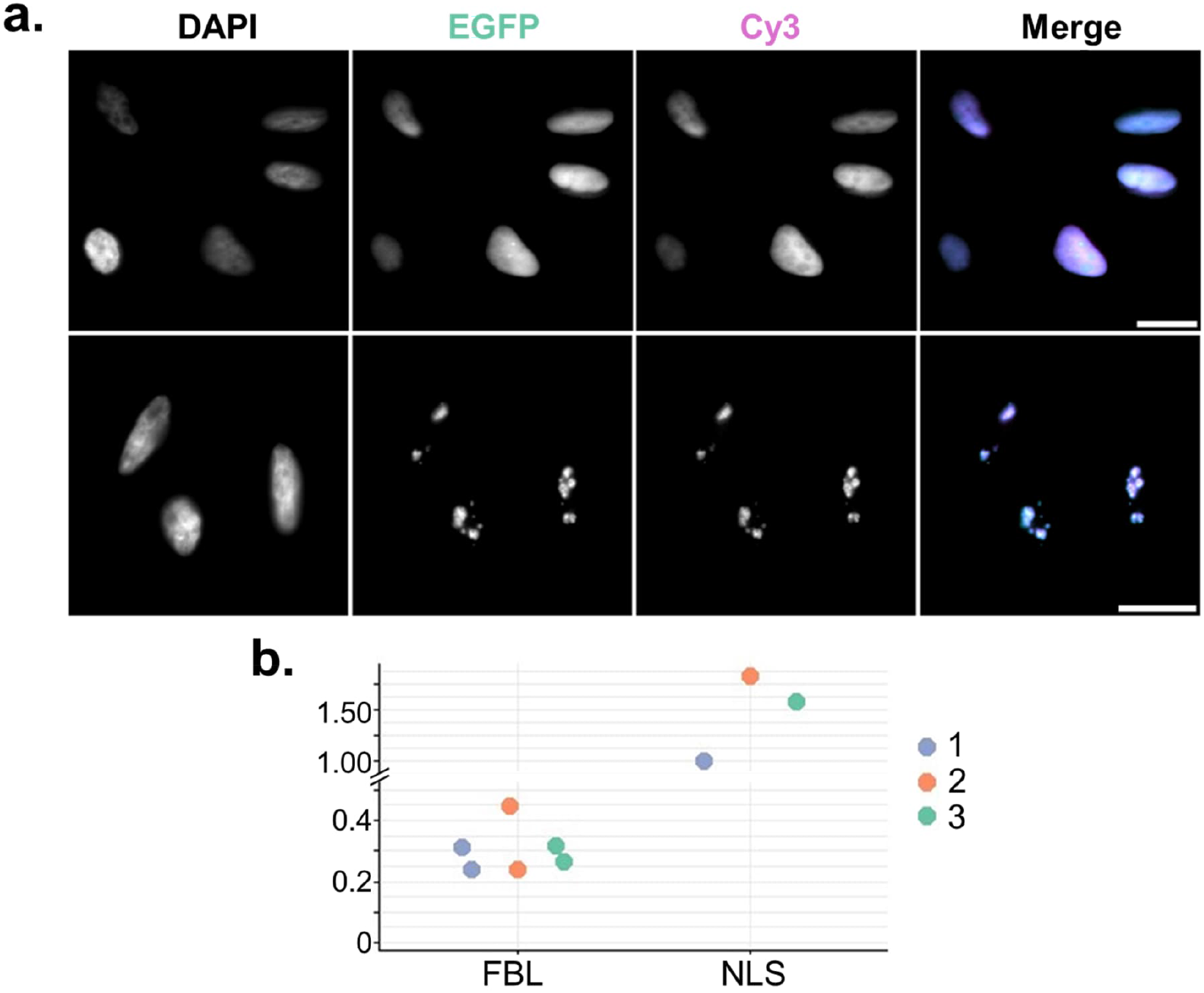
**a)** Fluorescent imaging of EGFP in the fusion protein and alkynylated RNA visualized by Cy3-azide. The colocalization of Cy3 and EGFP was used to evaluate the spatial specificity of the alkynylation reaction. The Halo-NLS and Halo-FBL fusion protein and the alkynylated RNA were restricted to the nucleus and nucleolus, respectively. Scale bar 20 µm. **b)** Percentual yields of the streptavidin pulldown from biotinylated input. For Halo-NLS samples, one streptavidin pulldown per replicate was performed, for Halo-FBL, two streptavidin pulldowns were pooled for each replicate.

Ribo-depleted libraries from the Halo-labeled RNA pulldown and input samples were sequenced using paired-end 150 bp reads, sequencing quality was evaluated with fastp. Only 0.5-1.6 % of reads were filtered out per sample, indicating high data quality. Salmon mapping yielded 28-47 million mapped reads per sample.

Following mapping and DEseq2 normalization, the reproducibility of Halo-seq samples was evaluated using Principal Component Analysis (PCA) (**Figure 3a**). Input and streptavidin-pulldown samples clustered along the first principal component (79% of variance), demonstrating high reproducibility across replicates. More than half of the significantly enriched and depleted RNAs in the pulldowns overlapped between the two Halo-fusions (**Figure 3b**). A substantial proportion of enriched RNAs were unique to each nuclear compartment (Halo-NLS vs Halo-FBL), indicating successful compartment-specific proximity labeling and RNA isolation.

**Figure 3.**
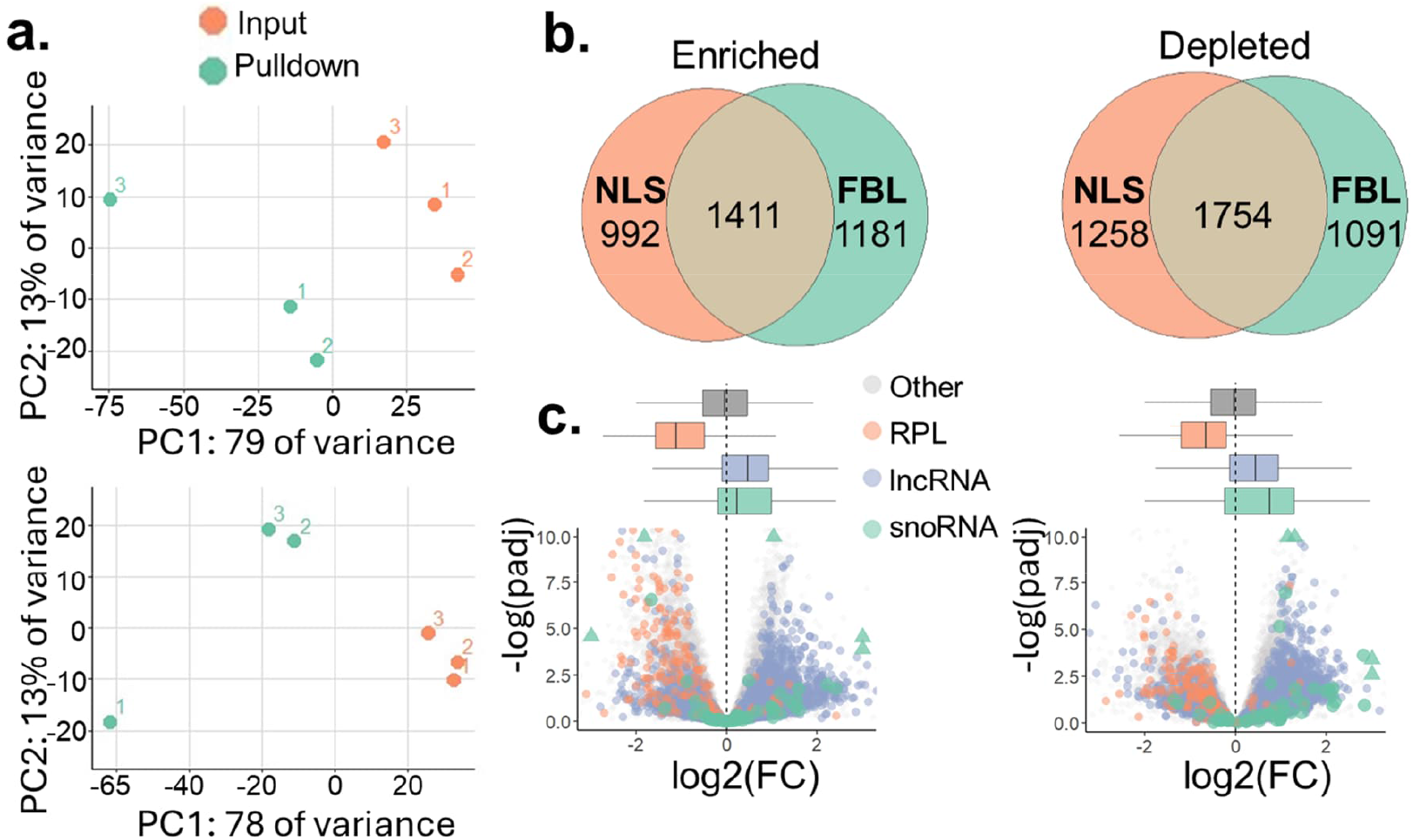
**a)** PCA-analysis of Halo-seq pulldown and input libraries. The samples separate on the first principal component according to experimental condition. **b)** Overlap of enriched and depleted genes in NLS and FBL Halo-seq pulldowns compared to their respective inputs. (enriched genes padj < 0.05 and LFC > 0.5; depleted genes padj < 0.05 and LFC < -0.5. **c)** Differential enrichment of RNA biotypes in Halo-seq pulldown and input samples. The volcano and box plots share the same x-axis. LncRNAs are enriched and ribosomal protein component encoding mRNAs (RPLs) depleted in the nuclear samples, snoRNAs are enriched in the nucleolus.

Distinct RNA biotypes displayed compartment specific enrichment and depletion patterns (**Figure 3c**), further confirming the spatial specificity of our Halo-seq. Cytosolic ribosomal protein mRNAs were depleted in both pulldowns, whereas nuclear localized lncRNAs were enriched in both compartments. Nucleolar snoRNAs were consistently enriched the Halo-FBL samples reflecting the nucleolar localization of Halo-FBL and the efficient labeling of RNAs in its proximity.

Taken together, these results demonstrate that our optimized Halo seq approach successfully isolates nuclear compartment specific RNA populations with high reproducibility and spatial precision.

Differences between RNAs captured in Halo-NLS and Halo-FBL proximity labeling were further examined by clustering genes into four groups: Cluster 1, enriched in both compartments; Cluster 2, enriched only in Halo-FBL pulldown; Cluster 3, enriched only in Halo-NLS pulldown, and Cluster 4, depleted in both compartments (**Figure 4a-b**). This clustering analysis further supported the spatial specificity of the proximity labeling approach.

**Figure 4.**
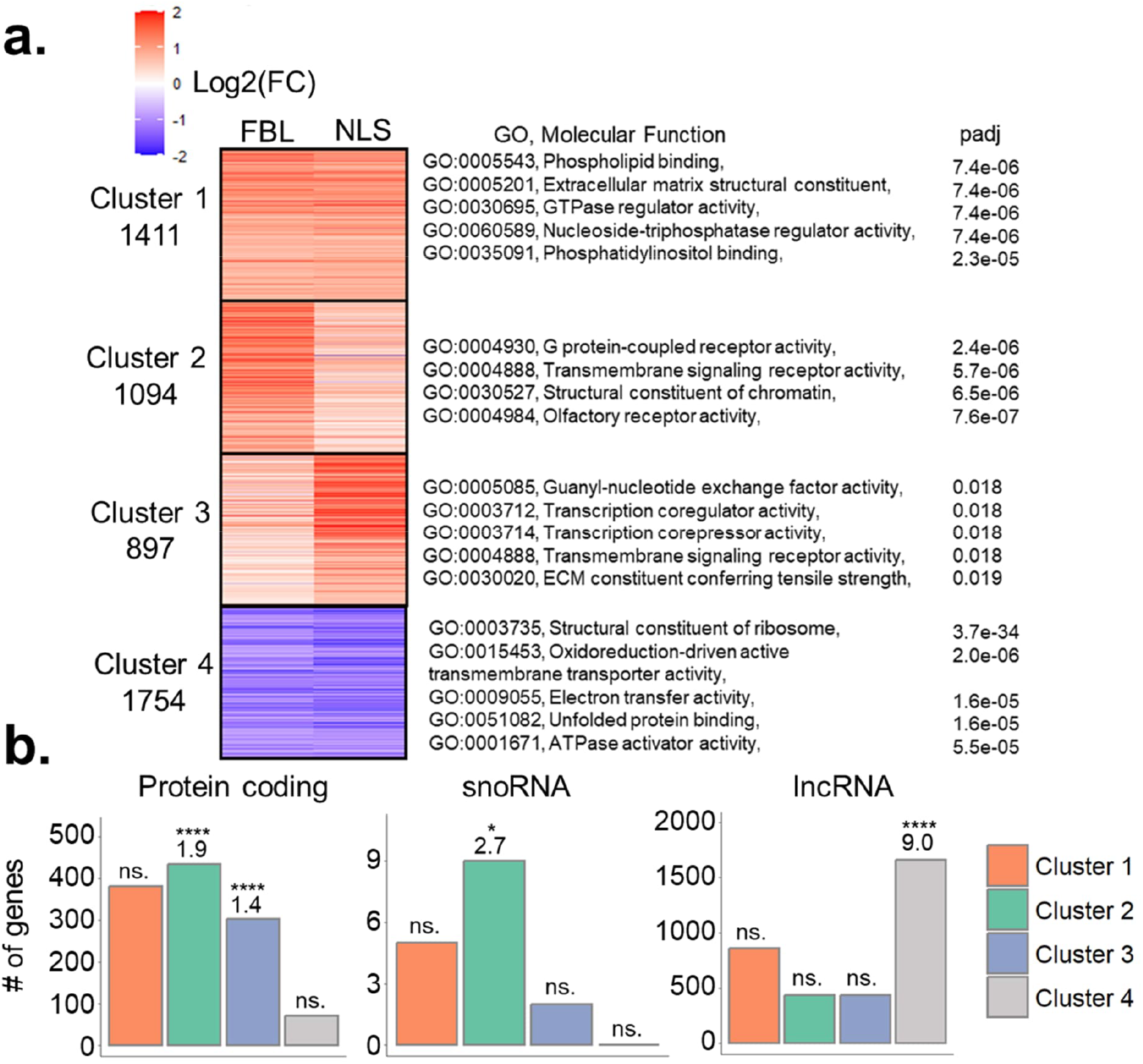
**a)** RNAs enriched/depleted in Halo-seq proximity labelling clustered based on the level of enrichment. The number of genes in each cluster is displayed on the left side of the heatmap. The right side displays the enriched molecular function GO-terms among protein coding RNAs in each cluster with the universe set to all expressed protein coding genes in the Halo-seq samples. **b)** RNAs in the four clusters shown by biotype. One-sided Fisher’s exact test was used to test overrepresentation of each biotype in the clusters. For significant p-values the odds ratio is shown.

Clusters 2 and 3 were significantly enriched for lncRNAs, consistent with their predominant localization within the nucleus. In contrast, Cluster 4 was enriched for protein-coding mRNAs, which are typically abundant in the cytosol and thus depleted in the nuclear compartments. Notably, snoRNAs were enriched exclusively in Cluster 2, reflecting the nucleolar specificity of Halo FBL and confirming successful extraction of nucleolar RNAs.

In conclusion, these analyses demonstrate that Halo-NLS and Halo-FBL proximity labeling effectively enrich nuclear RNAs with Halo FBL specifically capturing nucleolar RNAs—a subcompartment of the nucleus. Together, these results demonstrate the high quality of both our independently synthetized Halo-DBF ligand and the resulting Halo-seq proximity labeling dataset.

## Code Availability

No custom code was generated for the analyses in this manuscript. Code used for analysis and plotting is available upon request.

## Author Contributions

TA conducted experimental work and computational analyses. PT supervised the experimental work. MB assisted in the design of computational analyses. RL and AE synthesized the Halo-DBF ligand. PL made the plasmid constructs. DW designed and constructed the custom light-source. MLÄ and JK conceived the study and supervised experimental and computational work. TA and MLÄ compiled the manuscript.

## Acknowledgements

The authors acknowledge the Tampere University Imaging Facility and the Tampere University Flow Cytometry Facility.

## Data Availability

All raw fastq data files used in this study have been deposited to the Sequence Read Archive (SRA) under BioProject ID PRJNA1416425.

