## Supplemental text 1 for "Profiling of HeLa nucleoplasmic and nucleolar RNAs by Halo-seq proximity labeling"

### Profiling of HeLa nucleoplasmic and nucleolar RNAs by Halo-seq proximity labelling

Timo Ahola[1], Piia Takabe[1], Riku Salomäki[2], Päivi Lillsunde[1], Mohamed Bahrudeen[1], Donald Wlodkowic[3], Jan Kaslin[1,4,5], Alexander Efimov[2], and Minna-Liisa Änkö[1,*]

[1]Faculty of Medicine and Health Technology, Tampere University, Finland

[2]Faculty of Engineering and Natural Sciences, Tampere University, Finland

[3]School of Science, RMIT University, Melbourne, Australia

[4]Institute of Biomedicine, University of Turku, Finland

[5]Australian Regenerative Medicine Institute, Monash University, Australia

#### Synthesis of the DBF-ligand

4',5'-Dibromo-6-((2-(2-((6-chlorohexyl)oxy)ethoxy)ethyl)carbamoyl)-3-oxo-4a',9a'-dihydro-3H-spiro[isobenzofuran-1,9'-xanthene]-3',6'-diyl diacetate

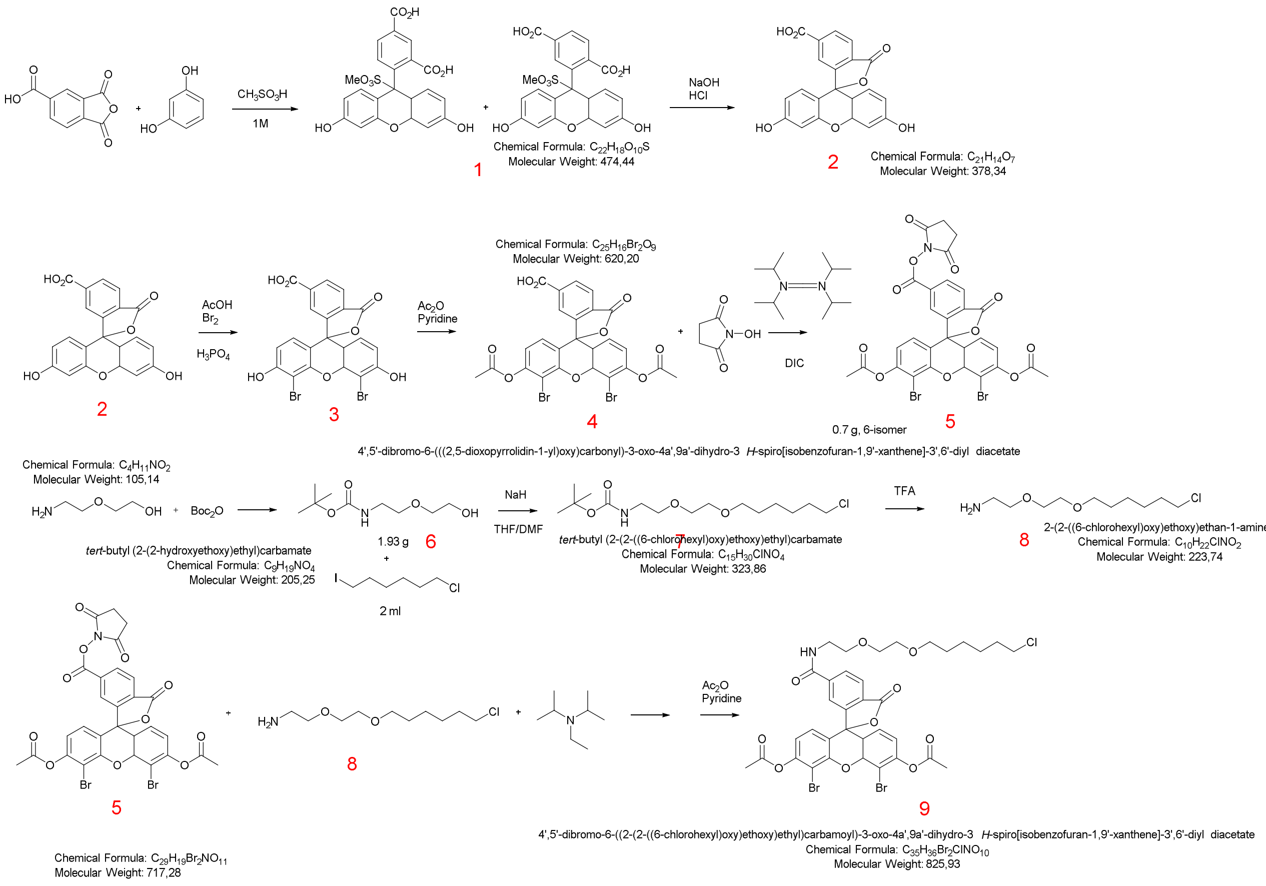


**General.** Solvents and reagents were obtained commercially from TCI Europe, Merck-Sigma-Aldrich-VWR and used as received unless otherwise noticed. Unless otherwise noticed, the glassware for the reactions was flame-dried and cooled under Ar in an exicator, and the reactions were performed under septa in the argon atmosphere. NMR spectra were recorded on a JEOL JNM-ECZ500R 500 MHz instrument and processed with JEOL Delta 5.2 software. Chemical shifts are reported in parts per million (ppm) and referenced to TMS (0.00 ppm) for $^{1}H$ NMR spectra and $CDCl_3$ (77.0 ppm) or DMSO-d6 (39.5 ppm) for $^{13}C$ NMR spectra. High-resolution mass spectra were measured on a JEOL TOF JMS-T100LP instrument with ESI+ or DART+ ionization using reserpine or PEG600 as a mass correction reference. The UV/vis absorption spectra were recorded with a Shimadzu UV-1800 spectrophotometer.

**Compound 1.** 2(4)-(3,6-dihydroxy-9-(methoxysulfonyl)-4a,9a-dihydro-9H-xanthen-9-yl)isophthalic acid. A total of 116 mg (1.05 mmol) of resorcinol, 101 mg (0.52 mmol) of trimellitic anhydride and 2 ml (31 mmol) of methane sulfonic acid were stirred in a 50 ml vial at 85-125 °C for 3 h. The solution was poured into 35 ml of ice water, the precipitate was filtered, washed and dried in vacuo at 150 °C to give 134 mg of **1** as a mixture of 2- and 4-carboxy isomers. NMR (DMSO-d6), δ ppm, $^{1}$H: 8.39 (s), 8.29 (d), 8.21 (d), 8.1 (d), 7.65 (s), 7.39 (d), 6.72 (m), 6.64 (m), 6.55 (m); $^{13}$C: 168.3, 166.5, 152.6, 152.5, 137.7, 136.6, 133.4, 131.5, 129.9, 127.4, 126.2, 125.3, 113.4, 109.7, 109.6, 102.8.

**Compound 2.** 3',6'-dihydroxy-3-oxo-4a',9a'-dihydro-3H-spiro[isobenzofuran-1,9'-xanthene]-5(6)-carboxylic acid. A total of 5.5 g of compound 1 was dissolved in 40 ml of 1M NaOH. Ca. 80 ml of 1M HCl was added slowly at stirring. A precipitate appeared immediately. The precipitate was filtered, washed with water, dried in vacuo to yield 4.2 g of the product **2**. NMR (DMSO-d6), δ ppm, $^{1}$H: 10.15 (s, 2H), 8.35-8.06 (m, 2H), 7.61-7.35 (m, 1H), 6.65-6.50 (m, 6H); $^{13}$C: 168.42, 168.4, 166.6, 160.2, 156.7, 152.4, 152.3, 137.8, 136.7, 133.3, 131.4, 130.0, 129.8, 129.7, 127.3, 126.0, 125.8, 125.1, 124.9, 113.2, 109.5, 109.4, 102.8, 83.9.

**Compound 3.** 4',5'-dibromo-3',6'-dihydroxy-3-oxo-4a',9a'-dihydro-3H-spiro[isobenzofuran-1,9'-xanthene]-5(6)-carboxylic acid. A total of 143 mg of NaOH were added to 1.5 ml of water and 1 g of ice, and stirred on ice bath for 10 minutes, after which 70 µl of bromine was added to the cold solution. The bromine solution was left stirring on ice bath until further use.

A total of 180 mg (0.48 mmol) of 2(4)-carboxy fluorescein **2** was suspended in 5 ml of water with 130 µl of EtOH. A 100 µl portion of 50%-weight NaOH/$H_2O$ was added to the solution, and the pH was estimated with an indicator paper. The pH was finally adjusted to be within 9.5-10.5 by adding a small portion of diluted NaOH.

The alkali bromine solution was added portion wise over the 30 min period to the fluorescein solution at stirring on cold water bath. The reaction mixture was stirred for another 30 minutes at r. t., after which the product was precipitated out by adding 279 µl of 85% phosphoric acid. The orange precipitate was filtered, washed with water and dried *in vacuo*, yielding 216 mg (84%) of the compound **3** as a mixture of 5(6)-carboxy isomers. HRMS (ESI-TOF) m/z 532.88541 found, calculated for open form $C_{21}H_{12}Br_2O_7$ 532.8870 $(M+H)^+$. NMR (DMSO-d6), δ ppm, $^{1}$H: 8.45-8.05 (m, 2H), 7.82-7.41 (m, 1H), 6.87-6.55 (m, 4H); $^{13}$C: 168.42, 168.4, 166.6, 160.2, 156.7, 152.4, 152.3, 137.8, 136.7, 133.3, 131.4, 130.0, 129.8, 129.7, 127.3, 126.0, 125.8, 125.1, 124.9, 113.2, 109.5, 109.4, 102.8, 83.9.

**Compound 4.** 3',6'-diacetoxy-4',5'-dibromo-3-oxo-4a',9a'-dihydro-3H-spiro[isobenzofuran-1,9'-xanthene]-5(6)-carboxylic acid. *Pyridine was dried by heating at reflux over NaOH for 1.5 h followed by a fractional distillation under Ar. The glassware for the reaction was dried at 160 °C for 2 h and cooled down under Ar.* A total fo 152 (0.28 mmol) of dibromofluorescein 3 was loaded into a 50 ml vial with containing 1.35 ml (14.28 mmol) of acetic anhydride and 35 µl (0.43 mmol, 1.54 eq) of dry pyridine. The solution quickly became pale yellow and was stirred at 80 °C for 3.75 h. The reaction mixture was concentrated on rotary evaporator, the residue was redissolved in dichloromethane and evaporated again. The dissolution-evapoartion was repeated for six times, after which the dry residue was dissolved in 20 ml of ethyl acetate and washed twice with 30 ml of saturated ammonium chloride. The organic phase was filtered through magnesium sulfate and evaporated. The solid was dried *in vacuo* to yield 158 mg of the compound **4** as a mixture of two isomers (TLC in DCM/EtOH 18/1). HRMS (ESI-TOF) m/z 616.90302 found, calculated for $C_{25}H_{14}Br_2O_9$ 616.90773 $(M+H)^+$. NMR ($CDCl_3$), δ ppm, $^1$H: 8.83-7.28 (m, 3H), 7.00-6.74 (m, 4H), 2.38 (s, 6H); $^{13}$C: 168.1, 152.5, 150.8, 149.1, 148.9, 136.1, 132.3, 129.9, 127.0, 126.9, 126.4, 126.0, 125.9, 119.7, 117.2, 106.9, 82.8, 21.2, 20.9.

**Compound 5.** 4',5'-dibromo-6-(((2,5-dioxopyrrolidin-1-yl)oxy)carbonyl)-3-oxo-4a',9a'-dihydro-3H-spiro[isobenzofuran-1,9'-xanthene]-3',6'-diyl diacetate. A flame-dried 40 ml vial with a stirring bar was charged with 102 mg (0.16 mmol) of the compound **4** and 0.75 ml of dry $CH_2Cl_2$. A 25 mg portion of N-hydroxysuccinimide and 30 µl of N,N'-diisopropylcarbodiimide were added, and the walls of the vial were rinsed with 2×0.75 ml of $CH_2Cl_2$. The vial was flushed with argon, sealed and left at stirring overnight.

The solvent was evaporated, and the solid residue was separated on flash column with 75 ml of Silica 60. Elution was done first with 480 ml of toluene:ethyl acetate 50:1 and yielded some pre-fractions; further elution with 350 ml of toluene:ethyl acetate 4:1 yielded the 5-isomer, an further elution with 150 ml of toluene:ethyl acetate yielded 110 mg of the target 6-substituted isomer. HRMS (ESI-TOF) m/z 735.90622 found, calculated for $C_{29}H_{17}Br_2NO_{11}Na$ 735.90606 $(M+Na)^+$. NMR ($CDCl_3$), δ ppm, $^1$H: 8.45-8.39 (m, 1H), 8.22-8.17 (m, 1H), 7.96 (s, 1H), 7.41-7.12 (m, H), 7.00-6.93 (m, 2H), 6.84-6.76 (m, 2H), 2.39 (s, 6H); $^{13}$C: 168.8, 168.0, 167.1, 160.4, 152.4, 150.9, 149.1, 132.6, 131.9, 131.1, 129.1, 129.0, 128.7, 128.3, 127.1, 126.4, 126.3, 125.4, 119.9, 116.9, 106.9, 82.0, 25.7, 20.9.

**Compound 6.** *tert*-butyl (2-(2-hydroxyethoxy)ethyl)carbamate. A flame-dried 40 ml vial with a stirring bar was charged with 945 µl (9.52 mmol) of 2-(2-aminoethoxy)ethanol and 20 ml of absolute ethanol, sealed and cooled down to 0 °C under argon. Boc anhydride (2.197 g, 10 mmol) was added, and the reaction mixture was stirred for 3 h. Solvents were evaporated, the dry residue was dissolved in 20 ml of dichloromethane and filtered though a pad of sodium sulfate. The solution was evaporated to yield 1.8 g of the compound **6** as an oil. HRMS (DART-TOF) m/z 206.13742 found, calculated for $C_9H_{19}NO_4$ 206.13868 $(M+H)^+$. NMR ($CDCl_3$), δ ppm, $^1$H: 5.36 (s, 1H), 3.78-3.67 (m, 2H), 3.62-3.50 (m, 4H), 3.39-3.08 (m, 3H), 1.45 (s, 9H); $^{13}$C: 156.3, 79.3, 72.3, 70.4, 61.6, 40.4, 28.5.

**Compound 7.** *tert*-butyl (2-(2-((6-chlorohexyl)oxy)ethoxy)ethyl)carbamate. A flame-dried 40 ml vial with a stirring bar was charged with 1.78 g of the compound **6**, 12.5 ml of dry THF and 6.3 ml of dry DMF. The vial was sealed, flushed with argon and cooled down to 0 °C on an ice bath. A total of 515 mg of NaH (60%) in mineral oil was added and the mixture was stirred on cold under Ar for 30 min. A portion of 1.8 ml of 6-chloro-1-idohexane was added, and the mixture was stirred at r.t. overnight. After 16 h the reaction was quenched with 40 ml of saturated solution of $NH_4Cl$, extracted with 40 ml of EtOAc, washed with water and brine, and the organic solvent was evaporated to give a yellowish oil. The TLC control with ninhydrine in EtOAc:hexane 3:7 revealed one major and a few smaller spots.

The compound was purified on 70 ml of Silica 60, eluted first with 280 ml of EtOAc:hexane 1:4, then with 350 ml of EtOAc:hexane 1:4. The second fraction contained the target compound, yield 990 mg (35%). HRMS (ESI-TOF) m/z 346.17864 found, calculated for $C_{15}H_{30}ClNO_4Na$ 346.17556 $(M+Na)^+$. NMR ($CDCl_3$), δ ppm, $^1H$: 5.00 (s, 1H), 3.64-3.36 (m, 10H), 3.35-3.17 (m, 2H), 1.80-1.68 (m, 2H), 1.45-1.26 (m, 13H); $^{13}C$: 156.0, 79.2, 71.4, 70.3, 70.1, 45.1, 40.4, 32.6, 29.5, 28.5, 26.8, 25.5.

**Compound 8.** 2-(2-((6-chlorohexyl)oxy)ethoxy)ethan-1-amine. A 40 ml flame-dried vial was charged with 420 mg of the compound **7** and 2 ml of dry dichloromethane. The solution was cooled down to 0 °C on ice bath under argon, and 0.5 ml of TFA was added. The reaction mixture was stirred on ice bath for 30 minutes, then at r.t. for 2 hours. Solvents were evaporated, the residue was dissolved in 15 ml of EtOAc and washed twice with saturated $NaHCO_3$ solution, dried over MgSO4. The organic solvent was evaporated in vacuo, yielding 240 mg (35%) of the product. The compound **8** was used in the next step without purification.

**Compound 9.** 4',5'-dibromo-6-((2-(2-((6-chlorohexyl)oxy)ethoxy)ethyl)carbamoyl)-3-oxo-4a',9a'-dihydro-3H-spiro[isobenzofuran-1,9'-xanthene]-3',6'-diyl diacetate. A flame-dried 40 ml vial equipped with a stirring bar was charged with 54 mg of the compound **5**, 60 mg of the compound **8** and 5 ml of dry dichloromethane. An aliquot of 60 µl of N,N-diisiopropylethylamine was added, the vial was sealed, flushed with argon and stirred at r.t. for 20 hours. The solvent was evaporated, the dry residue was redissolved in 6 ml of acetic anhydride and 1.5 ml of dry pyridine, and the solution was stirred in a dry sealed vial under argon at 80 °C for 3 hours.

The reaction mixture was evaporated in vacuo, the dry residue was redissolved in dichloromethane and evaporated again. The dissolution-evaporation in DCM was repeated five times, after which the dry residue was dissolved in 15 ml of EtOAc, washed with saturated solution of $NH_4Cl$, dried trough a pad of $MgSO_4$ and evaporated to dryness.

The solid residue was dissolved in 0.5 ml of chloroform and separated on four Silica 40 TLC plates 10×20 cm using the eluent hexane:EtOAc:MeOH 10:10:1 for development. The red spot with Rf 0.22 was collected to yield 45 mg (75%) of the target compound. HRMS (ESI-

TOF) m/z 844.01300 found, calculated for $C_{35}H_{34}Br_2ClNO_{10}Na$ 844.01302 $(M+Na)^+$. NMR ($CDCl_3$), δ ppm, $^1H$: 8.16-7.98 (m, 2H), 7.57 (s, 1H), 7.03-6.75 (m, 4H), 3.76-3.27 (m, 11H), 2.39 (s, 6H), 1.83-1.67 (m, 2H), 1.53-1.17 (m, 8H); $^{13}C$: 168.1, 168.0, 165.5, 152.9. 150.1, 148.9, 141.7, 129.7, 127.7, 125.9, 122.7, 119.7, 117.5, 106.7, 81.5, 71.2, 70.2, 70.0, 69.5, 45.1, 40.2, 32.5, 29.4, 26.7, 25.4, 20.9.
